# Signatures of vibration frequency tuning in human neocortex

**DOI:** 10.1101/2021.10.03.462923

**Authors:** Lingyan Wang, Jeffrey M. Yau

## Abstract

The spectral content of vibrations produced in the skin conveys essential information about textures and underlies sensing through hand-held tools. Humans can perceive and discriminate vibration frequency, yet the central representation of this fundamental feature is unknown. Using fMRI, we discovered that cortical responses are tuned for vibration frequency. Voxel tuning was biased in a manner that reflects perceptual sensitivity and the response profile of the Pacinian afferent system. These results imply the existence of tuned populations that may encode naturalistic vibrations according to their constituent spectra.

## INTRODUCTION

Our physical interactions with the environment produce complex, high frequency (>85Hz) vibrations in the skin whose spectral content underlie the manual perception of surface textures (Bensmaia & Hollins, 2005; Manfredi et al., 2014) and support sensing through hand-held tools (Brisben, Hsiao, & Johnson, 1999; Miller et al., 2018). Vibration frequency, like sound pitch, is a fundamental feature that we perceive and discriminate (Bolanowski, Gescheider, Verrillo, & Checkosky, 1988; Convento, Rahman, & Yau, 2018; Mountcastle, Talbot, Sakata, & Hyvärinen, 1969). Yet evidence for frequency-tuned somatosensory circuits remains conspicuously absent, in stark contrast to the tuning observed throughout the auditory neuraxis (Hudspeth, 2014; Saal, Wang, & Bensmaia, 2016; Wang, 2007). In human and non-human primates, vibration frequency is encoded in the periodicity of spiking activity of untuned cells in the peripheral afferent system (Johansson, Landstrom, & Lundstrom, 1982; Talbot, Darian-Smith, Kornhuber, & Mountcastle, 1968) and the earliest cortical processing stages (Harvey, Saal, Dammann 3rd, & Bensmaia, 2013; Lebedev & Nelson, 1996; Mountcastle et al., 1969). Conceivably, this temporal coding of vibration frequency gives rise to a rate-based representation in tuned populations, as seen in the auditory system (Saal et al., 2016; Wang, 2007). However, frequency-tuned somatosensory neurons have never been reported in primates and tuned cells were only recently discovered in the mouse somatosensory cortex (Prsa, Morandell, Cuenu, & Huber, 2019). The failure to establish frequency tuning in the primate brain may have been due to limited sampling of cortical territories or restricted exploration of vibrotactile stimulus space.

To search for vibration frequency tuning in the human brain, we performed whole brain functional magnetic resonance imaging (fMRI) as participants experienced a battery of vibrations on their hands while engaging in an attention-demanding frequency monitoring task (**Supplementary Fig. 1**). Vibrations, which were matched in perceived intensity, varied in frequency from 100 to 340Hz (**Supplementary Fig. 2**). We characterized voxel-level responses which reveal systematic tuning for vibration frequency. We compared voxel-tuning properties across participants and observed consistent tuning preferences that mirrored perceptual sensitivity and the response profile of the Pacinian afferent system. Lastly, we implemented an encoding model to provide an account for how voxel-level frequency tuning can relate to neural population responses.

## RESULTS

We first defined brain regions whose blood oxygen level-dependent (BOLD) activity was modulated by vibration stimulation applied to the left or right hands (**Fig. 1a**; **Supplementary Fig. 3**) irrespective of vibration frequency. Response modulation associated with right hand stimulation was greater in strength (*t*(6) = 2.48, *P* = 0.048) and more prevalent (*t*(6) = 4.21, *P* = 0.0056) compared to left hand responses. In both hemispheres of each participant, voxel responses were significantly modulated by vibrations delivered to the contralateral or ipsilateral hands (*F*-statistic: contralateral: 7.67 ± 0.86; ipsilateral: 7.37 ± 0.75). Response modulation associated with the contralateral and ipsilateral hands was similar in strength (*t*(6) = 2.28, *P* = 0.063) and prevalence (*t*(6) = 1.12, *P* = 0.30).

**Figure 1.**
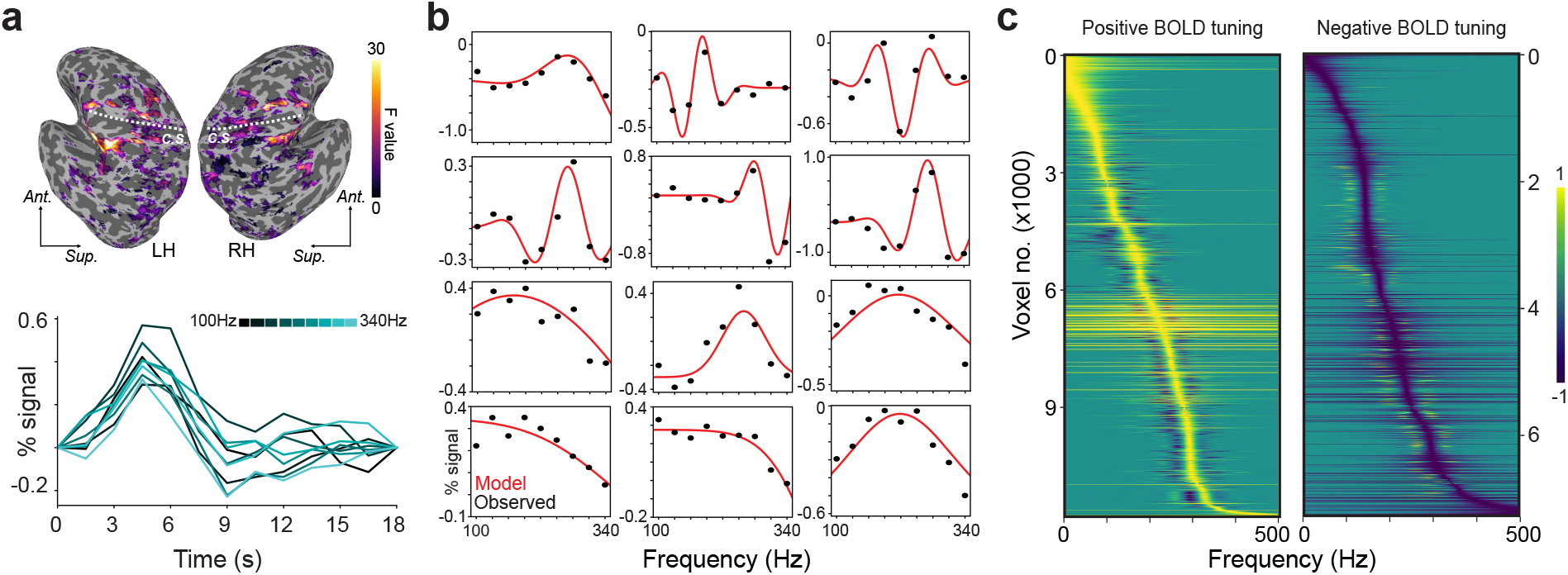
Frequency tuning of cortical responses to vibrations. (**a**) Vibrations delivered to the left or right hand are associated with significant BOLD signal modulation in an example participant's sensorimotor cortex. Dashed line indicates the central sulcus (c.s.) in the left hemisphere (LH) and right hemisphere (RH). BOLD signal time courses of an example voxel to different vibration frequencies follow stereotypical hemodynamic response profiles. Ant, anterior; Sup, superior. (**b**) Frequency tuning curves of example voxels (black dots indicate observed responses; red traces indicate fitted tuning functions). Frequency response patterns are consistent with Gabor tuning (top rows) or Gaussian tuning (bottom rows). (**c**) Normalized tuning curves for all frequency-selective voxels in example participant sorted by best modulating frequency (BF). Positive BOLD tuning voxels (left) exhibit signal increases at the BF while negative BOLD tuning voxels (right) exhibit signal decreases at the BF.

To characterize frequency-dependent modulation in vibration-responsive voxels (**Fig. 1b**), we fitted voxel-level response profiles with tuning functions (Materials and Methods). Tuning along a single dimension like temporal frequency can be modeled by fitting responses with simple Gaussian filters that parameterize the best modulating frequency (BF) and tuning sharpness. More complex frequency preferences can be modeled using Gabor filters that capture tuning profiles characterized by multiple modulation fields. Across participants, 59 ± 6.3% of vibration-responsive voxels exhibited significant tuning (FDR-corrected q < 0.05) that was described by the Gaussian model (*r* = 0.67 ± 0.013; range: 0.23-0.97) or Gabor model (*r* = 0.57 ± 0.025; range: 0.36-0.95). Tuned voxels were predominantly found in parietal and frontal cortex (**Supplementary Fig. 4**). We performed model selection for each tuned voxel (Materials and Methods) and found that voxel profiles tended to be more consistent with Gaussian tuning rather than Gabor tuning (proportion of voxels consistent with Gabor tuning = 0.40 ± 0.12). We compared the prevalence of Gaussian vs Gabor tuning in different sensorimotor regions under the assumption that simpler tuning could define primary sensory areas while more complex tuning could be confined to higher-order areas. Across regions, we observed similar proportions of tuned responses best described by the Gaussian and Gabor models (**Supplementary Fig. 5**). That voxels characterized by Gaussian- and Gabor-shaped tuning are interspersed in parietal and frontal brain regions is inconsistent with the notion that simple frequency selectivity gives way to more complex tuning over a somatosensory cortical hierarchy.

The number of tuned voxels in the left hemisphere (5685 ± 3353) and right hemisphere (5961 ± 3635) did not differ significantly (*t*(6) = 1.35, *P* = 0.23) (**Supplementary Table 1**). While most tuned voxels were selective for only one hand, voxels tuned to contralateral and ipsilateral stimulation were observed in both hemispheres, and 20.59 ± 6.06% of tuned voxels were selective for vibrations applied to either hand (**Supplementary Table 2**). Because contralateral and ipsilateral stimulation has been associated with BOLD signal increases and decreases (Schäfer et al., 2012), respectively, we tested whether tuned voxels were more likely to exhibit negative BOLD signal changes with ipsilateral stimulation. Voxels exhibited signal increases and decreases (**Fig. 1b**), but the likelihood for tuned voxels to deactivate at their BF did not differ between contralateral and ipsilateral stimulation across participants (*t*(6) = 1.08, *P* = 0.32) or within each participant (z-statistic = −0.028–0.23, *P* = 0.81–0.99). For tuned voxels with positive or negative activity changes, frequency response profiles spanned the entire range of tested frequencies (**Fig. 1c**). These results imply the existence of cortical feature detectors that are selective for the frequency components comprising naturalistic vibrations (Manfredi et al., 2014).

Having established voxel-level frequency tuning, we asked whether voxel preferences were biased to frequencies near 250Hz, the range corresponding to the maximum response sensitivity of the Pacinian afferent system (Bell, Bolanowski, & Holmes, 1994; Bolanowski & Verrillo, 1982; Johansson et al., 1982) and the peak perceptual sensitivity in humans (Bolanowski et al., 1988; Bolanowski & Verrillo, 1982). In each participant, BF distributions (**Fig. 2a**) differed significantly from uniform (1-sample Kolmogoroz-Smirnov test; all *P* < 1e-15) with more voxels preferring intermediate frequencies (BF: 222 ± 22 Hz; range: 187-258 Hz) compared to lower and higher frequencies (**Supplementary Table 3**). We additionally tested whether voxels tuned for both hands had similar frequency preferences for the left and right hands (**Fig. 2b**), but BF values were uncorrelated between hands (*r* = −0.021 ± 0.088; *t*(6) = - 0.58, *P* = 0.58). The finding that cortical frequency representations, which are maintained independently for the left and right hands, mirror the sensitivity profile of human observers and of the peripheral afferent system is consistent with efficient coding theory (Barlow, 1961).

**Figure 2.**
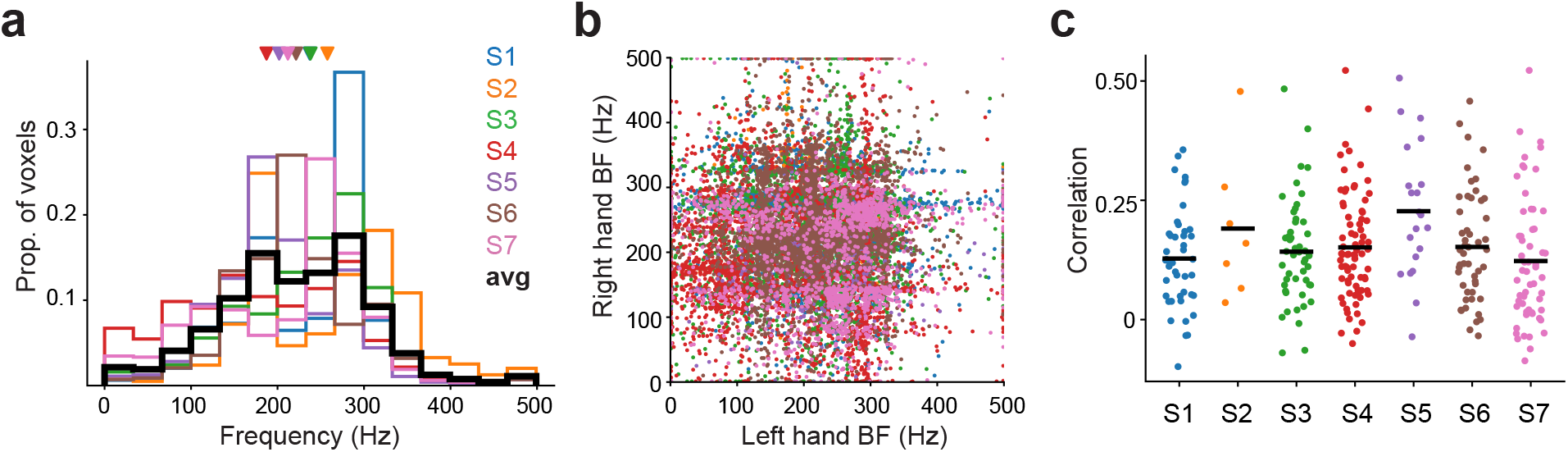
Best modulating frequency distributions within participants, between hands, and across activation clusters. (**a**) Distribution of best modulating frequency (BF) in each participant (N = 7). Colors indicate individual participants. Average distribution is denoted in black. Triangles indicate mean BF in each participant. (**b**) Relationship between left hand BF and right hand BF in voxels tuned for both hands. Frequency preferences were uncorrelated over hands (mean r = −0.021 ± 0.088; t(6) = −0.58, P = 0.58). (**c**) Relationship between voxel locations and frequency preferences. Dots indicate correlation between the physical distances separating voxel pairs within an activation cluster and their BF differences. The average correlation for each participant is denoted by the black bar. Although correlations were generally positive, BF maps were unstructured and inconsistent with tonotopic organization.

In the auditory cortical system, the spatial clustering of neurons with similar frequency preferences produces orderly tonotopic maps that are resolvable with fMRI (Barton, Venezia, Saberi, Hickok, & Brewer, 2012; Martino et al., 2015). We wondered whether an analogous topography, based on vibration frequency tuning, exists in the somatosensory cortical system. We first tested if the spatial proximity between pairs of frequency-tuned voxels within activation clusters related to the similarity of their frequency preferences (Materials and Methods). The physical distances between voxels were correlated with their BF differences (**Fig. 2c**) for left hand responses (*r*: 0.18 ± 0.038; *t*(6) = 11.30, *P* = 2.87e-5) and right hand responses (*r*: 0.14 ± 0.042; *t*(6) = 8.41, *P* = 0.00015), implying that voxels with similar preferences tended to aggregate. This aggregation alone, however, is insufficient evidence for tonotopic organization because neighboring voxels could share frequency preferences simply due to spatial smoothing effects or the point-spread function of BOLD (Shmuel, Yacoub, Chaimow, Logothetis, & Ugurbil, 2007). Indeed, BF maps in each participant were generally disordered and lacked global structure (**Supplementary Fig. 4**). We further evaluated BF maps using a more conservative tonotopy definition that assumed frequency preferences within an activation cluster would be arranged in a gradient over the cortical surface (Materials and Methods). A mere 0.60% of the total activation clusters (2 out of 336 over all participants) comprised voxels with BFs spanning the full frequency range that were arranged in a gradient. The weak evidence for orderly tonotopic maps implies that frequency tuning does not define somatosensory cortical topography.

Because participants were all right-hand dominant (Edinburgh handedness scores: 87 ± 3.6) and they selectively attended to vibrations delivered to their right hands during the scans, we reasoned that response profiles may differ between hands. Such differences would presumably be reflected in the distributions of tuning model parameters, which were highly consistent across participants (**Supplementary Fig. 6**). Indeed, we observed greater response modulation with right hand responses (**Supplementary Fig. 6a**) (*t*(6) = 2.47, *P* = 0.048), although baseline activity levels were equivalent over the hands (**Supplementary Fig. 6b**) (*t*(6) = 1.63, *P* = 0.16). We predicted that right hand responses would be more frequency selective, but tuning widths did not differ between the hands (**Supplementary Fig. 6c**) (*t*(6) = 1.94, *P* = 0.10). For voxels best described by the Gabor model, we evaluated phase parameter distributions and found that phase distributions differed between hands in all participants (Watson's two-sample test of homogeneity; *U*^2^ = 0.82–5.66, *P* < 0.001). Despite these differences, phase distributions were typically bimodal (Rayleigh test, *P* < 1e-15) with prominent peaks at 0.5π and 1.5π that indicate a general tendency for tuning functions to comprise balanced positive and negative peaks (**Supplementary Fig. 6d**). Altogether, these analyses highlight the consistency of tuning patterns across participants and reveal differences between left and right hand tuning profiles that may be related to hand dominance or attention.

How might voxel-level tuning be related to cortical population activity? Vibrations delivered to the glabrous skin entrains the activity in some cortical populations and frequency could be represented by a spike timing code using these untuned but phase-locking neurons (Harvey et al., 2013; Lebedev & Nelson, 1996; Mountcastle et al., 1969). However, the frequency-response profiles of these neurons – characterized by spike rates that increase monotonically with frequency – are incompatible with voxel-level Gaussian and Gabor tuning, assuming the BOLD signal reflects aggregate population activity (Klink, Chen, Vanduffel, & Roelfsema, n.d.).

Alternatively, vibration frequency could be carried in the activity of tuned populations, which have recently been identified in mouse somatosensory cortex (Prsa et al., 2019). Phase-locking responses are less prominent as one ascends the cortical hierarchy (Harvey et al., 2013), which may reflect a transition to a rate-based code. As a proof of concept, we implemented an encoding model to explore how the activity of tuned populations could relate to voxel-level responses (Materials and Methods). We assumed that a voxel's response reflects the weighted combination of activity in neural populations selective for different frequencies (**Fig. 3a**). These encoding models recapitulated observed voxel profiles (**Fig. 3b**) and accounted for substantial response variance (**Fig. 3c;** scaled goodness-of-fit: 0.68 ± 0.086). We also verified that the encoding models captured voxel tuning by performing a decoding analysis (Materials and Methods). The models predicted multivoxel activity patterns that closely resembled observed patterns (**Fig. 3d**). Accordingly, a simple decoder (**Fig. 3e**) identified the frequencies associated with different measured patterns with an accuracy (67% ± 23%) far exceeding chance performance (11%) (*t*(6) = 5.82, *P* < 0.0011). Lastly, we considered how phase-locking populations could contribute to voxel responses and found that voxel tuning could be recapitulated only if the phase-locking neurons exhibited some degree of frequency selectivity (**Supplementary Fig. 7**). These modeling results confirm that frequency information is carried in voxel responses and provide a conceptual framework for relating voxel tuning to frequency-selective neural populations.

**Figure 3.**
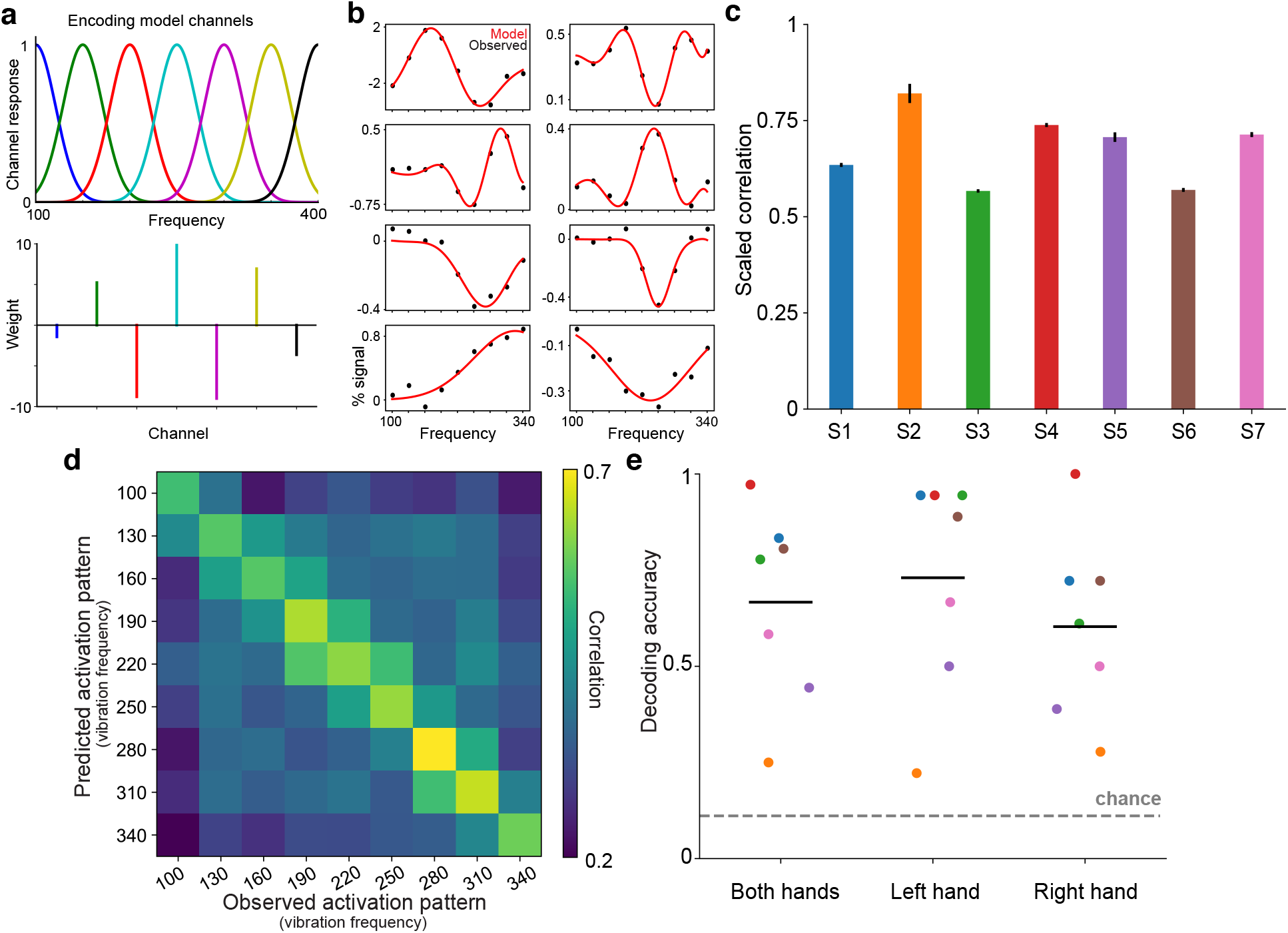
Encoding model based on activity of putative frequency-tuned neural populations. (**a**) Gaussian channels represent neural populations that respond selectively to different vibration frequencies. The encoding model assumes that a voxel's response to any given vibration frequency is the weighted sum of the activity in the channels. (**b**) Encoding model captures tuned response patterns in example voxels (black dots). Red curves indicate model-predicted responses profiles. (**c**) Bars indicate voxel-averaged scaled model performance within each participant. The model is trained on one fold of data and tested on a held-out fold. Model performance is the correlation between the model predictions and the test data, normalized by the correlation between the two folds of data (which represent the maximum correlation possible given the noise in the data). Error bar indicate s.e.m. (**d**) Correlation matrix indicates the similarity between multivoxel activation patterns predicted by the encoding model and observed patterns in the held-out data. Correlations are averaged over hands and participants. For decoding, an algorithm identifies the model-predicted pattern yielding the highest correlation with an observed pattern to infer the frequency condition. (**e**) Cross-validated decoding performance for both hands and each hand separately. Black line indicates group averaged accuracy. Colored dots indicate individual participants. Dashed line indicates chance performance.

## DISCUSSION

We find that voxel-level BOLD signals are modulated by vibrations in a manner that clearly reflects frequency selectivity. Voxel tuning spans the range of frequencies that are relevant for fine texture perception (Bensmaia & Hollins, 2005; Manfredi et al., 2014). Frequency preferences are consistent across individuals and systematic with a greater share of voxels preferring frequencies that optimally drive the Pacinian afferent system (Bell et al., 1994; Bolanowski & Verrillo, 1982; Johansson et al., 1982). This cortical bias may underlie our enhanced perceptual sensitivity for vibrations near 250Hz (Bolanowski et al., 1988; Bolanowski & Verrillo, 1982). Our finding that cortical representations mirror environmental statistics, peripheral afferent profiles, and perceptual sensitivity is consistent with the predictions of efficient coding theory (Barlow, 1961).

Conceivably, frequency-tuned voxel activity reflects neurons in the primate somatosensory cortical system that are analogous to vibration selective cells recently identified in mice. Importantly, although individual neurons in mouse somatosensory cortex exhibit vibration tuning (Prsa et al., 2019), such neurons have never been reported in the primate brain. Limited sampling of cortical populations and territories may have obscured the presence of frequency-tuned neurons. Alternatively, frequency tuning may be a property that only emerges at a population level in primates. Arbitrating between these possibilities will require large scale neurophysiological recordings, which can be guided by our neuroimaging findings.

Future studies will also need to address the mechanisms that generate frequency selectivity in the somatosensory system: Our data reveal frequency selective cortical responses despite the absence of fine tuning in peripheral and subcortical processing stages. This contrasts with the auditory system, where frequency tuning exists throughout the neuraxis, even at the receptor level (Hudspeth, 2014). Cortical tuning may reflect the central convergence of submodality signals that are initially carried by distinct populations in the peripheral afferent system (Pei, Denchev, Hsiao, Craig, & Bensmaia, 2009; Saal & Bensmaia, 2014; Saal, Harvey, & Bensmaia, 2015). In fact, recent evidence has challenged the traditional functional dichotomy between Pacinian and non-Pacinian perceptual channels by positing a universal frequency decoding system (Birznieks et al., 2019). Beyond submodality convergence, varying distributions of excitatory and inhibitory neurons may also underlie the diversity of frequency-selective population responses across sensory cortex (Hughes et al., 2021). At a cellular level, short term synaptic depression may impose a frequency dependent filter on information transmission (Rosenbaum, Rubin, & Doiron, 2012) and mediate the conversion from temporal coding to rate coding (Lee, Wang, & Bendor, 2020).

Regardless of the mechanism, our data reveal somatosensory cortical activity in human neocortex that is tuned for vibration frequency. Analogous frequency encoding schemes in the somatosensory and auditory systems may facilitate the extensive crosstalk between touch and audition in the temporal frequency domain (Crommett, Madala, & Yau, 2019; Crommett, Perez-Bellido, & Yau, 2017; Yau, Olenczak, Dammann, & Bensmaia, 2009; Yau, Weber, & Bensmaia, 2010). Moreover, frequency-selective cortical filters offer an efficient scheme for representing the complex spectra of vibrations encountered in naturalistic touch.

## MATERIALS AND METHODS

### Participants

Seven healthy adult volunteers (5 females; mean age ± SD: 26 ± 2.8 years; aged 20-29 years) participated in the study. All participants were right-handed (Oldfield, 1971) (Laterality quotient: 87.1 ± 9.7). The sample size was set on the basis that any significant and consistent outcomes established in 7 out of 7 subjects would be statistically generalizable according to a 2-tail binomial test (*P* < 0.05). Participants had normal or corrected-to-normal vision. Testing procedures were approved by the Baylor College of Medicine Institutional Review Board. All participants provided written consent and were paid for their participation or waived payment.

### MRI acquisition

All scans were conducted in the Core for Advanced MRI (CAMRI) at Baylor College of Medicine. MRI data were acquired on a 3-Tesla MAGNETOM Trio scanner with Prisma fit (Siemens, Erlangen, Germany) using a 64-channel head coil. Anatomical data were acquired using a T1-weighted magnetization prepared rapid acquisition gradient echo sequence (MPRAGE; TR = 2300 ms; TE = 2.98 ms; flip angle = 9°; 1 mm^3^ voxels). Functional data were obtained using an axial echo-planar imaging (EPI) sequence with simultaneous multi-slice (SMS) acceleration (TR = 1500 ms; TE = 33 ms; flip angle = 90°; GRAPPA factor = 2; SMS factor = 3; FOV = 192 mm; 69 slices; 2 mm^3^ voxels; 380 volumes per scan) that covered all of the cortical volume and part of the cerebellum. Each participant underwent 12 functional scans (~9.5min/scan) divided across 2 sessions (5.9 +/- 7.4 days inter-session interval).

### Tactile stimulation

Vibrotactile cues were delivered to the distal pad of the participant's left and right index fingers using an MRI-compatible piezoelectric tactor (Engineering Acoustics, Inc., Casselbery, FL). Tactors were fastened to the distal finger pads with self-adherent cohesive wrap bandages. Tactors were controlled using the EAI Tactor Development Kit and stimulus timing was determined using custom Matlab scripts. The vibration set comprised 9 frequencies: 100, 130, 160, 190, 220, 250, 280, 310, and 340Hz. Vibrations were matched in perceived intensity with amplitudes (gain: 71.4–97.4 arbitrary units according to EAI controller) determined in preliminary behavioral experiments using the method of adjustment. To further ensure that participants attended to vibration frequency rather than intensity during the scans, we applied a random ±5% jitter in amplitude on each stimulus presentation. Offline, we measured vibration amplitudes (unloaded) using a laser displacement sensor (ZX2-LD50, Omron, Hoffman Estates, IL) (displacement range: 0.414–0.504mm) and confirmed tactor reliability (**Supplementary Fig. 2**).

### Frequency monitoring task and scans

Participants were scanned in an event-related design as they performed a vibration frequency monitoring (oddball detection) task (Perez-Bellido, Barnes, Crommett, & Yau, 2017) while maintaining visual fixation. Each scan comprised unimanual and bimanual events. An event comprised a series of 3 vibration stimuli (stimulus duration: 700ms; inter-stimulus interval: 300ms). On the majority of events (*regular events*; 66/76 in each scan corresponding to 2 repetitions each of 9 right hand frequencies, 9 left hand frequencies, and 15 bimanual frequency combinations), the frequency of the three vibrations was identical. All of the analyses included in this report were based on the unimanual regular events. On a subset of events (*oddball events*; 10/76 in each scan), the frequency of the second vibration differed from the first and third vibrations in the series (frequency difference: 120-240Hz). Participants were instructed to report the occurrence of oddball events using a foot pedal response (Current Designs, Philadelphia, PA). Reliable detection of oddball events (**Supplementary Fig. 1**) indicated that participants attended to vibration frequency. The responding foot was counter-balanced across sessions over participants. To control for spatial attention effects, oddball events only occurred on the right hand (on unimanual and bimanual events) such that attention was directed toward each subject's dominant hand throughout the scan. Events were separated by 3, 4.5, 6, or 7.5-s intervals with order and timing determined pseudo-randomly using Optseq2 (http://surfer.nmr.mgh.harvard.edu/optseq).

### Behavioral analysis

Given the unequal number of oddball and regular events, we quantified oddball detection performance by computing an F_1_ score for each subject (Powers, 2011):

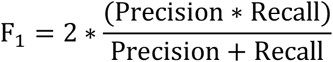

where *Precision* is defined as the number of hits (correctly detected oddball events) divided by the sum of hits and false alarms (events incorrectly identified as oddball) and *Recall* is defined as the number of hits divided by the sum of hits and misses (oddball events not detected). F_1_ scores range from 0–1 with higher scores indicating better performance. For each subject, we determined if the observed F_1_ score was greater than that expected by chance (**Supplementary Fig. 1**) by generating a null distribution of F_1_ scores assuming the observed number of positive responses with shuffled event labels over 1000 permutations.

### fMRI analyses

Data preprocessing and first-level analyses were performed using AFNI (Cox, 1996). Each participant's data were preprocessed using standard procedures (*afni_proc.py*) including: (i) slice timing correction (*3dTshift*); (ii) image co-registration (*align_epi_anat.py*); (iii) functional image alignment (3dvolreg); (iv) spatial blurring with a 4-mm FWHM filter (3dmerge); (v) mean-normalization of each voxel's signal (*3dcalc*). Preprocessed voxel-wise data were modeled using multiple linear regression (*3dDeconvolv*e): general linear models (GLM) comprised 34 regressors corresponding to left hand stimulation (9 frequencies), right hand stimulation (9 frequencies), 15 bimanual conditions, and oddballs. Each regressor was created using a gamma-variate convolution kernel. The GLM comprised head motion and drift parameters as nuisance regressors. GLM coefficients were taken as the voxel response associated with each condition. A single GLM was fitted to the whole 12-scan dataset to define the analysis mask comprising voxels whose activity was modulated by either left hand or right hand stimulation. For the tuning models and encoding model analyses, separate GLMs were fitted after dividing the full dataset into 2 folds corresponding to the 6 scans from each scanning session. Unless otherwise noted, analyses were performed in native volume space. For displaying purposes, each participant's data were projected into surface space. Surface models were constructed from each participant's anatomical scans using Freesurfer (Dale, Fischl, & Sereno, 1999). The analysis exploring the topographic organization of frequency preferences was performed in native surface space.

In each participant, we defined an analysis mask by identifying voxels whose activity was modulated by either left hand or right hand stimulation. For each hand separately, an omnibus F-statistic was computed to quantify the significance of each voxel's responses to the 9 vibration frequencies. The full analysis mask was the union of the left hand and right hand F-statistic maps, thresholded at a false discovery rate (FDR) corrected *q* < 1e-4 over the whole brain.

### Vibration frequency tuning functions

To test for vibration frequency tuning, we fitted parametric tuning functions to each voxel's frequency response profiles estimated from the 2 data folds. If a voxel's response profiles were inconsistent over the folds, a tuning model fit to these data would be meaningless. Accordingly, we only fit tuning functions to voxels whose across-fold Pearson correlation exceeded 0.2. For voxels with consistent profiles across folds, we fitted simple and complex tuning functions and performed model competition to determine the model most appropriate for each voxel given its complexity and performance. Models were fitted to each voxel's response data using the method of least squares which minimized the error between observed and predicted data. Responses to left and right hand stimulation were considered separately.

To capture simple frequency tuning, we assumed a voxel's response profile was characterized by a Gaussian function:

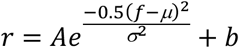

where *r* is the predicted voxel response to a vibration with frequency *f*, *A* is a gain term, *μ* is the best modulating frequency, *σ* is the tuning width, and *b* indicates the baseline activity level over all frequencies. The Gaussian model comprised 4 free parameters.

To capture more complex frequency tuning patterns, we assumed a voxel's response profile was characterized by a Gabor function, a cosine wave modulated by a Gaussian window:

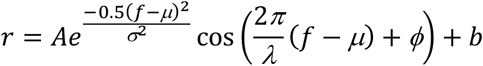

where *r* is the predicted voxel response to a vibration with frequency *f*, *A* is a gain term, *μ* is the center of the Gaussian, *σ* is the spread of the Gaussian, *λ* and *ϕ* are the wavelength and phase of the wave, and *b* indicates the baseline activity level over all frequencies. The Gabor model comprised 6 free parameters. In preliminary analysis, we found that estimating *λ* with no constraints could yield small wavelength values that reflected the noise in the data. Accordingly, we constrained *λ* by requiring the *λ/σ* ratio to be >2.25 in the final analysis.

To determine whether a voxel’s responses were better captured by the Gaussian or Gabor models, we compared models using Akaike information criterion (AIC) (Burnham & Anderson, 2004):

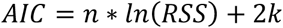

where *n* is the number of data points used to fit the models, *RSS* is the residual sum of squared errors, and *k* is the number of free parameters. The AIC-preferred model of each voxel was taken as that which yielded the smaller AIC value. We then computed the correlation between the AIC-preferred model predictions and the observed data to quantify goodness-of-fit. Voxels were considered to be tuned if the correlation between model predictions and observed data was statistically significant after correcting for the number of modeled voxels (FDR corrected *q* < 0.05 using the Benjamini-Hochberg procedure).

We evaluated a number of features defining voxel tuning curves (**Supplementary Fig. 6**). We defined the peak of a tuning function as the curve portion corresponding to the greatest (modulus) response modulation. Best modulating frequency (BF; 1-500 Hz) was the *μ* parameter for Gaussian models or the frequency corresponding to the peak for Gabor models. The gain term indicated the (unsigned) magnitude of response modulation. The baseline parameter represented basal activity common to all frequencies. Tuning sign (positive or negative) corresponded to direction of activity change relative to the baseline level at the peak. The full width at half maximum (FWHM) along the peak indicated the tuning selectivity of each voxel.

### Voxel-wise encoding models

We implemented a simple channel encoding model to predict voxel-level activity by assuming the existence of frequency-tuned cortical neurons like those recently identified in mouse somatosensory cortex (Prsa et al., 2019). We modeled the normalized activity levels (*R*) of a cortical population in response to a vibration with frequency *f* using a Gaussian channel:

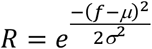

where μ is the population’s BF and σ is the channel tuning width. We assumed a voxel comprises different populations with unique frequency preferences, so the full encoding model predicted a voxel’s response (*r*) as a linear combination of activity from multiple populations:

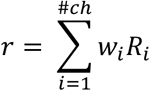

where *R*_*i*_ is the normalized activity of the *i*^*th*^ population and *w*_*i*_ is a weight that describes the population’s contribution to the voxel’s overall response. We modeled each voxel using 7 channels with predefined BF values. Accordingly, an encoding model was fitted to the response profiles of each tuned voxel by estimating the channel weights and a tuning width parameter that was shared over all the channels. Model fitting was performed using 2-fold cross-validation. Parameters were estimated using the method of least squares to minimize the error between model predictions and the tuning curve describing one data fold. Model performance was computed as the Pearson correlation between the model predictions and the data in the second fold. The final goodness-of-fit was the cross-validated model performance averaged over the two folds. Because the cross-validated goodness-of-fit depends on the consistency of the two folds, we normalized model performance by the across-fold correlation and report scaled correlations. For voxels tuned to both hands, separate models were fitted to explain left hand and right hand responses.

In separate analyses, we considered how voxel level activity may be related to phase-locking neurons that have been identified in non-human primates (Harvey et al., 2013; Lebedev & Nelson, 1996; Mountcastle et al., 1969) (**Supplementary Fig. 7**). We reasoned that the total spiking activity of these neurons would be minimal at low vibration frequencies and grow with increases in vibration frequency. Importantly, phase-locking neurons would fail to respond on every stimulus cycle at high vibration frequencies because of neural refractoriness, so population firing rates would saturate. We modeled this ramp-to-plateau response profile of a neural population using a rectified linear unit (ReLU) as a channel in our encoding model:

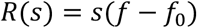

where *R(s)* is the normalized population activity to a vibration with frequency *f*, *s* is a slope parameter describing the relationship between population activity and frequency, and *f*_*0*_ is the lowest frequency at which the population responds (set to 1Hz). Because neural activity depends on vibration amplitude (Harvey et al., 2013) and populations can differ in their sensitivity to vibration amplitude, we modeled different populations (i.e., channels) as rectified linear units with different slopes. Note that by allowing the channels to have different slopes, we assume that neural populations in a voxel respond differentially over vibration frequencies thereby building frequency tuning into the model. The full encoding model, then, predicted a voxel’s response (*r*) as a linear combination of activity from multiple populations:

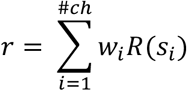

where *R*(*s*_*i*_) is the normalized activity of the *i*^*th*^ population defined by slope *s*_*i*_ and *w*_*i*_ is a weight that describes the population’s contribution to the voxel’s overall response. We assumed each voxel comprised 8 populations with predefined slopes. The ReLU models were trained and tested in the same manner as the Gaussian channel model.

We implemented a simple decoder using the encoding models to verify further that the models captured the frequency response profiles of the voxels. The decoding analysis included only the voxels with significant encoding model performance (*P* < 0.05). Using the encoding models fitted to one data fold, we generated multivoxel activity patterns for each vibration frequency. These patterns served as labeled templates against which the observed multivoxel activity patterns in the other data fold could be compared. For decoding, we computed the correlations between an observed activation pattern and each of the template patterns predicted with the encoding models. The template pattern yielding the maximum correlation was taken as the decoded frequency. For each participant, decoding performance was the accuracy averaged over the two folds. Because distinct voxel sets exhibited tuning for left and right hand stimulation, the encoding and decoding analyses were performed separately for each hand.

### Topography analysis

We performed two analyses to establish evidence for a topographic organization based on voxel frequency preferences. We first tested whether the physical distance (in volume space) between pairs of voxels related to the similarity of their BFs. For each participant, this analysis was performed within activation clusters (minimum cluster size = 40 voxels). For each cluster, we defined 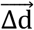 as a vector of distances between each pair of voxels and 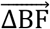 as a vector of pairwise voxel BF differences. We computed the correlation between 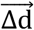 and 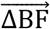 for each activation cluster. At the group level, we tested whether the average (within participant) correlation over clusters differed significantly from 0.

The second analysis tested whether frequency preferences within activation clusters were arranged in a gradient pattern over the cortical surface (minimum cluster size: 60 surface nodes). Two conditions needed to be met in order for an activation cluster to be considered tonotopic. First, the cluster needed to contain nodes with BFs that spanned the full frequency range. For each cluster, we binned BF values from 50–450 Hz in 50-Hz steps. We only further considered clusters that had at least one node in each BF bin. Second, BF values within a cluster were required to be systematically arranged. For each cluster, we defined an axis that passed through the cluster’s center. We then projected each node’s BF onto the axis and performed linear regression between the BF values and node locations along the axis. A significant linear regression fit indicated that BFs were ordered in a gradient along the axis. Because we were agnostic to the orientation of potential BF gradients, we defined repeated the analysis along 4 axes (0°, 45°, 90°, and 135°) for each cluster.

### Statistical analysis

Statistical analyses in this paper include Pearson correlation, pair-wise t test, one-sample t test, one-sample Kolmogorov-Smirnov uniformity test, and the two-sample independent proportions test. For circular data, we performed the Rayleigh uniformity test and Watson’s two-sample test of homogeneity. All tests were performed using Python 3.7 or R 3.5.1.

## ACKNOWLEDGEMENTS

This work was supported by R01NS097462 (JMY). We acknowledge BCM’s Core for Advanced MRI (CAMRI) and the Computational and Integrative Biomedical Research Center (CIBR). We thank Md. Shoaibur Rahman with data collection. We thank Nuo Li, Mike Beauchamp, Yue Zhang, and Meghan Robinson for helpful feedback and technical support. We thank Jing Lin for help with vibrometry and James Romesberg for assistance with constructing the foot pedal. We are grateful to Yau Lab members for helpful discussions.

## AUTHOR CONTRIBUTIONS

J.M.Y. designed the experiment. L.W. collected the data. L.W. and J.M.Y. wrote the analysis code, analyzed, and interpreted the data. L.W. and J.M.Y. wrote the manuscript.

## Supplementary Material

**Supplementary Fig. 1.**
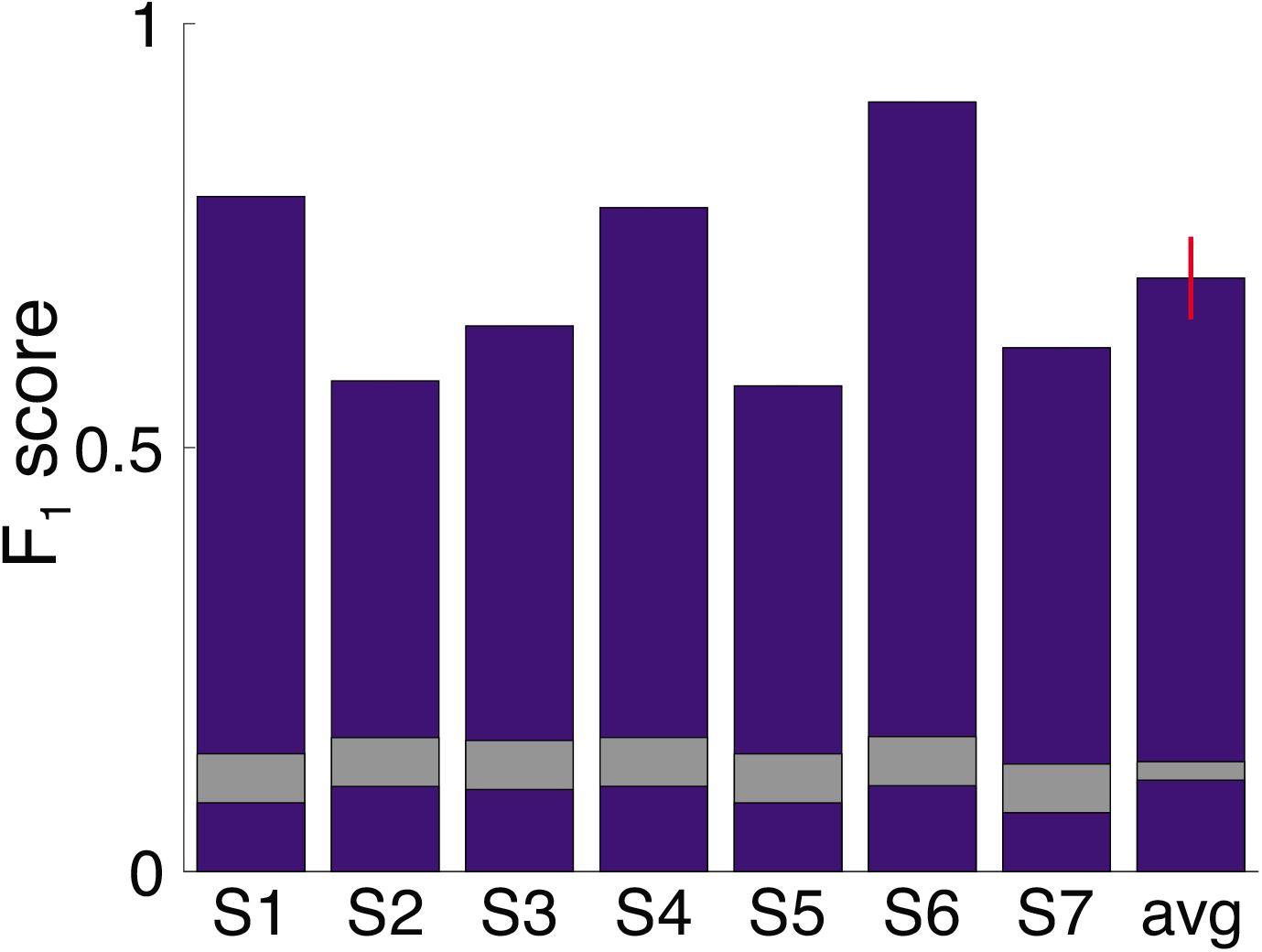
Oddball detection performance. F_1_ score indexes detection performance by accounting for precision and recall^1^. Higher scores indicate better performance. Bars indicate F1 score for each participant and the group-averaged score. Red error bar indicates s.e.m. Gray segments indicates the F_1_ score distributions expected by chance (center = mean score; thickness = standard deviation) given the number of positive responses provided by each participant (Materials and Methods). Performance in each participant far exceeded chance levels.

**Supplementary Fig. 2.**
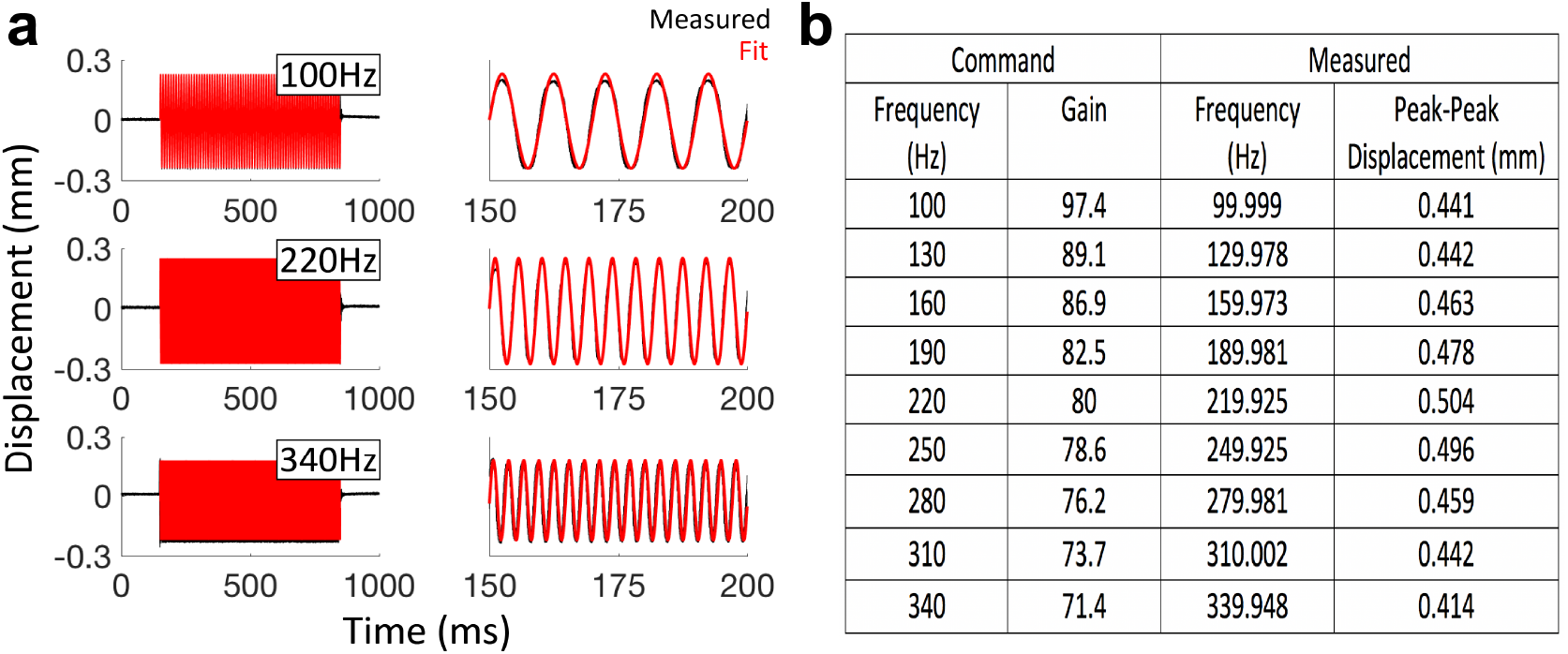
Analysis of measured displacement (**a**) Unloaded vibration amplitudes were measured outside of MRI environment using a ZX2-LD50 Laser Displacement Sensor (response time 240μs; acquired by Power 1401 CED with 40kHz sampling rate). Displacement profiles for 10 repeats (black) of the 100-, 220-, and 340-Hz stimuli are shown along with fitted sinusoids (red). Waveforms on the right show cycles from a portion of the full measurements. (**b**) Table indicates command frequencies and gains used to drive stimuli with the Engineering Acoustics, Inc (EAI) controller as well as the measured frequencies and displacements.

**Supplementary Fig. 3.**
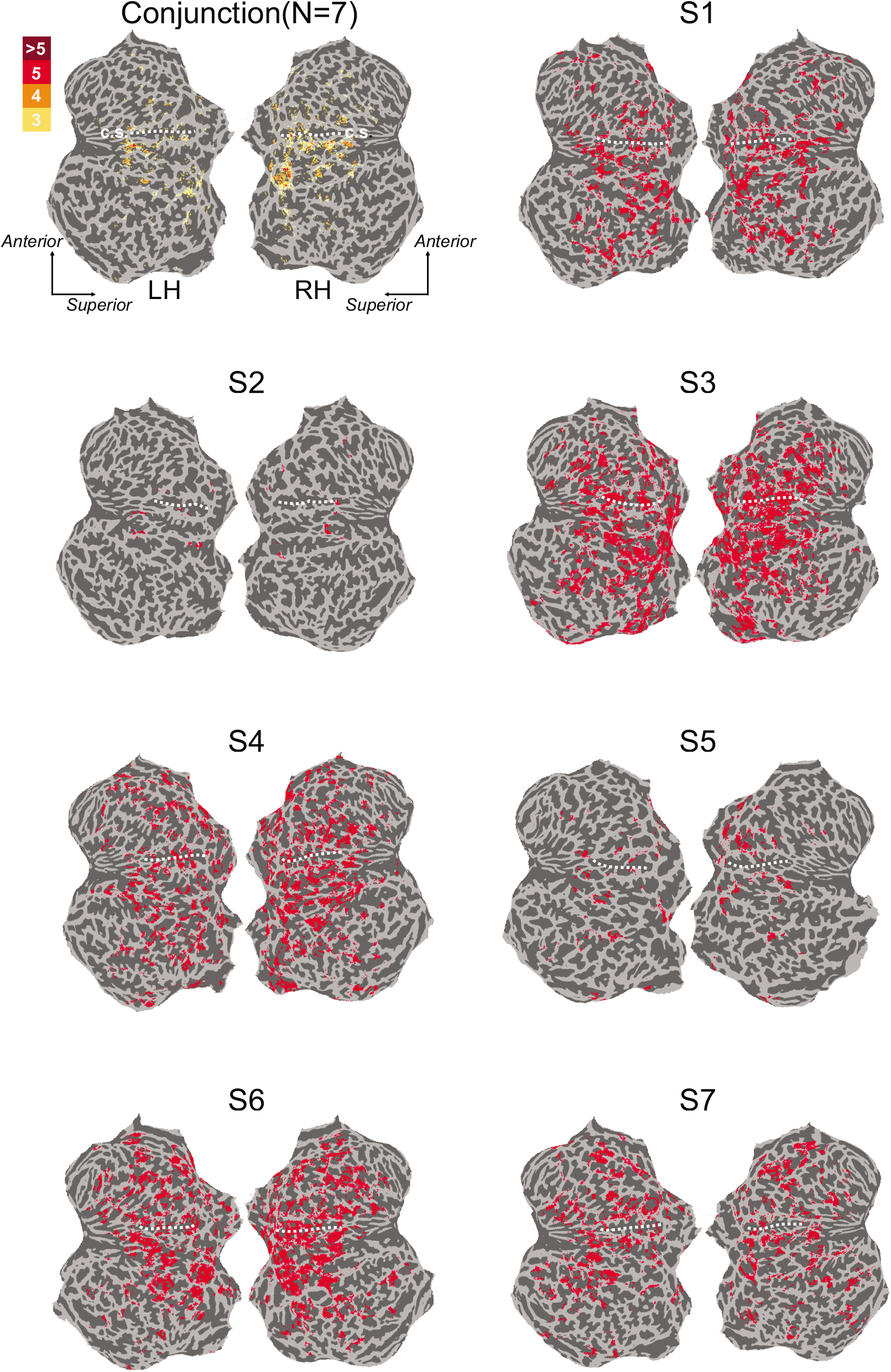
Analysis masks for each participant and group summary mask. Labeled surface nodes indicate significant responses to left or right hand vibrations (FDR corrected q = 0.0001). Conjunction map indicates nodes with significant activations in 3 or more participants. Dashed white lines indicate the central sulcus (c.s.). LH, left hemisphere; RH, right hemisphere

**Supplementary Fig. 4.**
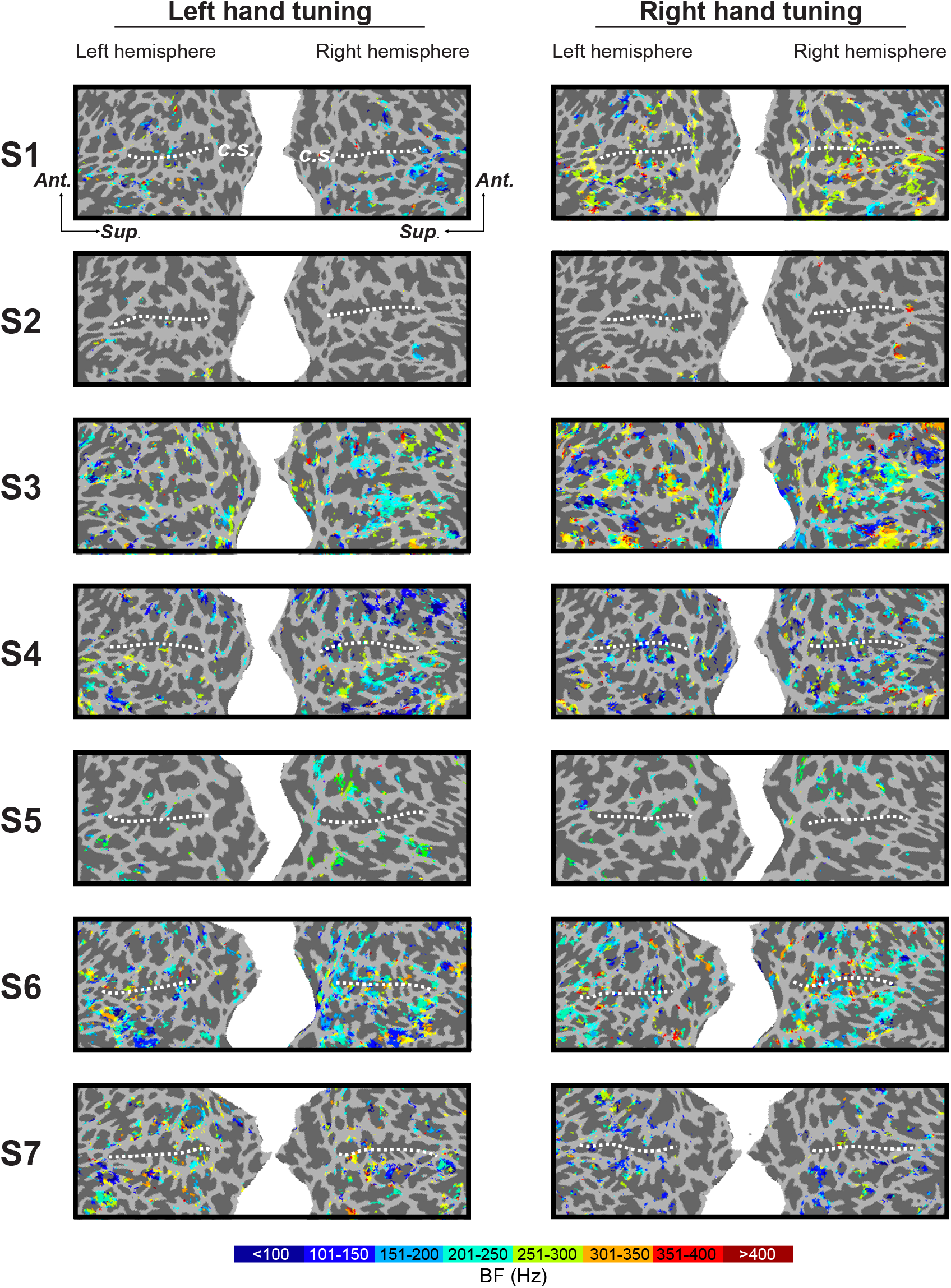
Frequency preference maps. Best modulating frequency (BF) for significantly tuned voxels in each participant are projected onto cortical surfaces. Left and right hand tuning is depicted in separate maps. Dashed white lines indicate the central sulcus (c.s.). Ant., anterior; Sup., superior

**Supplementary Fig. 5.**
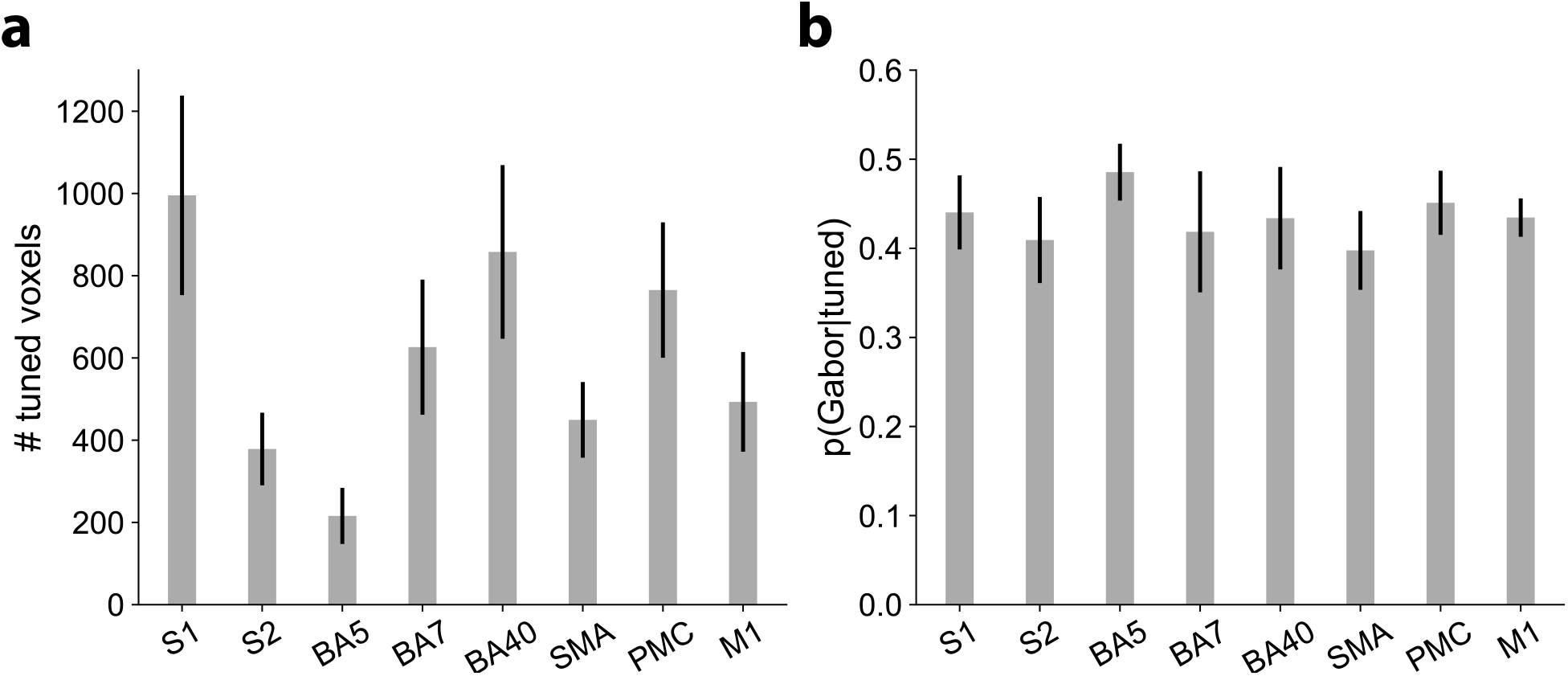
Voxel tuning across sensorimotor cortical regions. Regions are defined using Human Connectome Project parcellations^2^. (**a**) Number of tuned voxels in sensorimotor regions. (**b**) Proportion of tuned voxels in each sensorimotor region with response profiles more consistent with Gabor tuning rather than Gaussian tuning according to Akaike Information Criterion. BA, Brodmann area; S1, primary somatosensory cortex; S2, secondary somatosensory cortex; SMA, supplementary motor area; PMC, premotor cortex; M1, primary motor cortex.

**Supplementary Fig. 6.**
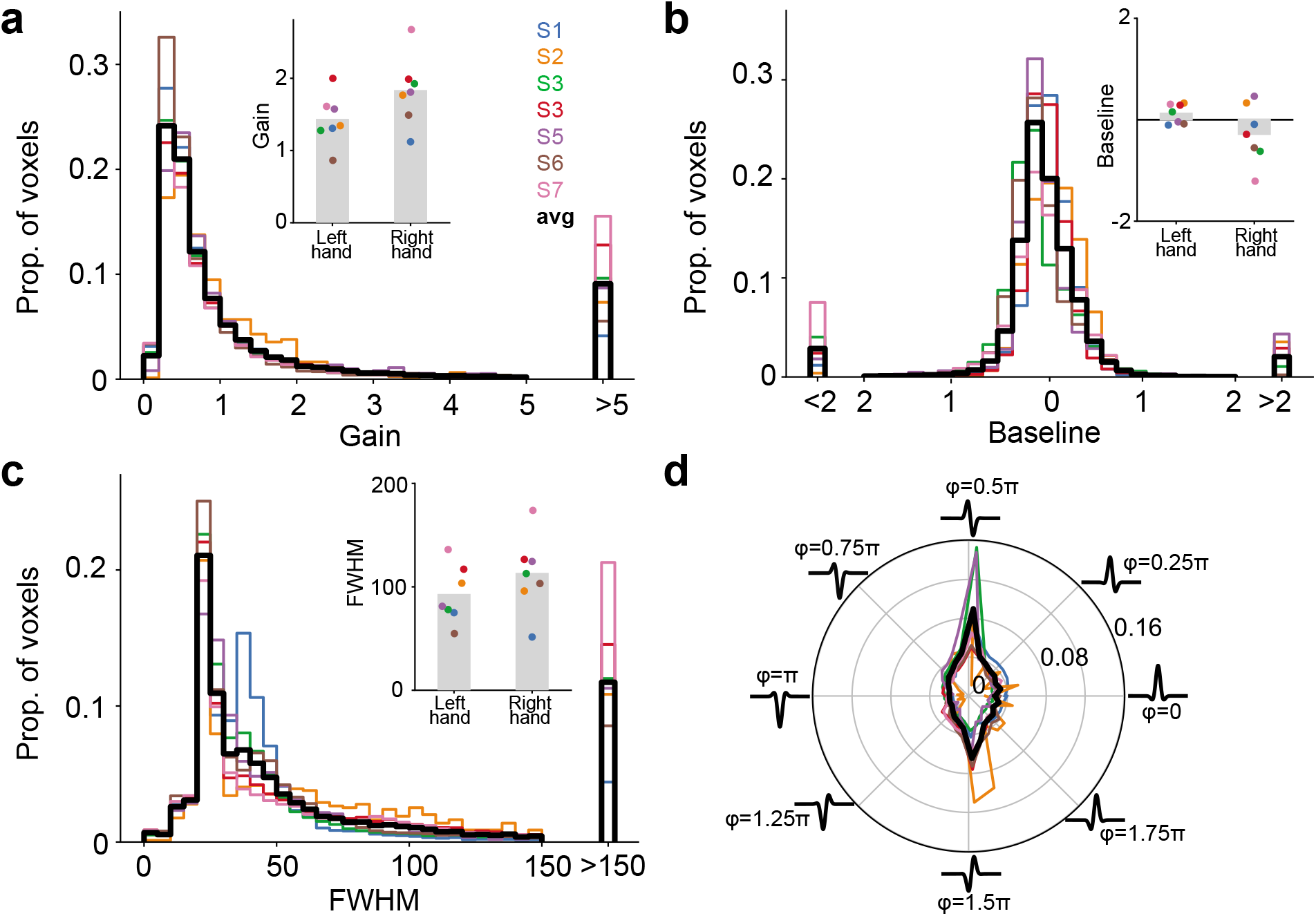
Frequency tuning parameters are highly consistent across participants. (**a**) Distribution of gain parameters from Gaussian and Gabor functions fitted to tuned voxels. Black traces indicates group average. Inset shows gain parameters sorted according to left hand and right hand tuning functions. Bars indicates group average and dots indicate individual participant averages. Gains were significantly larger for right hand tuning (*t*(6) = 2.47, *P* = 0.048). (**b**) Distribution of baseline parameters from Gaussian and Gabor functions fitted to tuned voxels. Conventions as in *a*. Baseline values did not differ significantly between hands (*t*(6) = 1.63, *P* = 0.16). (**c**) Distribution of frequency selectivity as indexed by the full-width at half-maximum (FWHM) of the each Gabor or Gaussian tuning function’s dominant peak. Conventions as in *a*. FWHM did not differ significantly between hands (*t*(6) = 1.94, *P* = 0.10). (**d**) Distribution of phase parameter values from the Gabor tuning functions. Plotted Gabors indicate canonical profile associated with each phase value. Conventions as in *a*. Although phase distributions differed between hands (Watson’s two-sample test of homogeneity; *U*^2^ = 0.82– 5.66, *P* < 0.001), there was a consistent pattern for non-uniform phase distributions (Rayleigh test, *P* < 1e-15) with peaks at φ = 0.5π and 1.5π.

**Supplementary Fig. 7.**
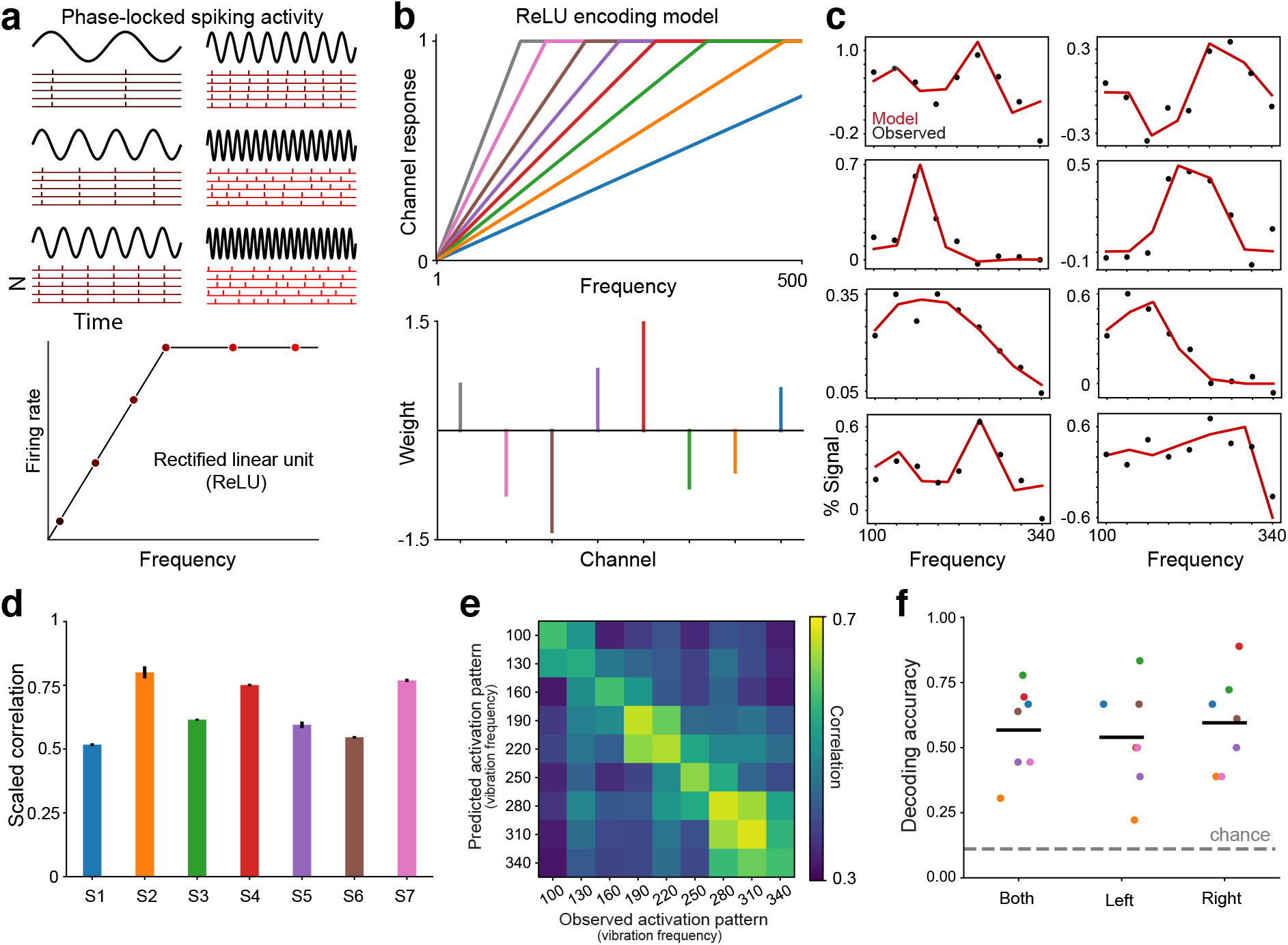
Encoding model based on activity of phase-locking neural populations. (**a**) Vibration frequency can be encoded in the timing of spiking activity in somatosensory cortical neurons^3–5^. Rasters show idealized activity in neurons whose spikes occur at particular phases of each vibration cycle. At low frequencies, the neurons can fire on every cycle. At high vibration frequencies, phase-locking neurons may skip cycles occurring during their refractory period. Accordingly, the frequency response profile for phase-locking neurons can be described by a low-frequency range over which rates increase monotonically before plateauing at higher frequencies. This profile is captured by a rectified linear activation unit (ReLU). (**b**) ReLU encoding model assumes that a voxel’s response to any given vibration frequency is the weighted sum of the activity in a set of ReLU functions (representing different populations) with different slopes. Slopes are hyperparameters and the weights are estimated in model fitting. Note that the assumption of different slopes implies that the neural populations represented by the ReLU functions are implicitly selective for frequencies. (**c**) ReLU encoding model (red trace) captures tuned response patterns in example voxels (black dots). (**d**) Bars indicate voxel-averaged scaled model performance within each participant. The model is trained on one fold of data and tested on a held-out fold. Model performance is the correlation between the model predictions and the test data, normalized by the correlation between the two folds of data (which represent the maximum correlation possible given the noise in the data). Error bars indicate s.e.m. (**e**) Correlation matrix indicates the similarity between multivoxel activation patterns predicted by the encoding model and the held-out fold. For decoding, the algorithm identifies the model-predicted pattern yielding the highest correlation with an observed pattern to infer the frequency condition. (**f**) Cross-validated decoding performance for both hands and each hand separately. Black line indicates group averaged accuracy. Colored dots indicate individual participants. Chance performance is 11%.

**Supplementary Table 1.** Number of tuned voxels in the left and right hemispheres.

| PARTICIPANT | LEFT | RIGHT |
| --- | --- | --- |
|  | HEMISPHERE | HEMISPHERE |
| 1 | 6009 | 6701 |
| 2 | 273 | 321 |
| 3 | 10357 | 11027 |
| 4 | 7668 | 8024 |
| 5 | 1423 | 1329 |
| 6 | 7974 | 8877 |
| 7 | 6088 | 5446 |

Counts indicate the number of tuned voxels in each participant. Vibration-responsive voxels were considered tuned only if they exhibited reliable across-fold correlations ( $r > 0.2$ ) and significant tuning function fits (FDR corrected $q < 0.05$ ). A voxel was included in the counts only once regardless of whether it was tuned for both hands.

**Supplementary Table 2.** Number of voxels tuned to the contralateral hand, ipsilateral hand, or both hands.

| PARTICIPANT | CONTRA | IPSI | BOTH |
| --- | --- | --- | --- |
| 1 | 5348 | 4986 | 2376 |
| 2 | 148 | 248 | 198 |
| 3 | 8684 | 9267 | 3433 |
| 4 | 6640 | 5646 | 3406 |
| 5 | 1167 | 1152 | 433 |
| 6 | 5917 | 6906 | 4028 |
| 7 | 5370 | 4467 | 1697 |

Counts indicate the number of voxels over the left and right hemispheres in each participant that exhibited tuning only for the contralateral (CONTRA) or the ipsilateral (IPSI) hand. Voxels that were tuned for both hands are indicated in the 3<sup>rd</sup> column.

**Supplementary Table 3.** Voxel-level frequency preferences.

| PARTICIPANT | BF MEAN | BF MEDIAN | KS STAT | P VALUE |
| --- | --- | --- | --- | --- |
| 1 | 237 | 267 | 0.99 | 1e-15 |
| 2 | 258 | 273 | 0.99 | 1e-15 |
| 3 | 238 | 252 | 0.99 | 1e-15 |
| 4 | 187 | 179 | 0.99 | 1e-15 |
| 5 | 202 | 195 | 0.99 | 1e-15 |
| 6 | 221 | 216 | 0.99 | 1e-15 |
| 7 | 212 | 238 | 0.99 | 1e-15 |

Values indicate best modulating frequency (BF) statistics in each participant. KS, Kolmogoroz-Smirnov test statistic

